# Testing soil nematode extraction efficiency using different variations of the Baermann funnel method

**DOI:** 10.1101/318691

**Authors:** Annika Eva Schulz, Nico Eisenhauer, Simone Cesarz

## Abstract

Nematodes are increasingly used as powerful bioindicators of soil food web composition and functioning in ecological studies. Todays’ ecological research aims to investigate not only local relationships but global patterns, which requires consistent methodology across locales. Thus, a common and easy extraction protocol of soil nematodes is needed. In this study, we present a detailed protocol of the Baermann funnel method and highlight how different soil pre-treatments and equipment (soil type, amount of soil, sieving, filter type) can affect extraction efficiency and community composition by using natural nematode communities. We found that highest nematode extraction efficiency was achieved using lowest soil weight (25 g instead of 50 g or 100 g) in combination with soil sieving, and by using milk filters (instead of paper towels). PCA at the family level revealed that different pre-treatments significantly affected nematode community composition. Increasing the amount of soil increased the proportion of larger-sized nematodes being able to overcome long distances. Sieving is suggested to break up soil aggregates and, therefore, facilitate moving in general. Interestingly, sieving did not negatively affect larger nematodes that are supposed to have a higher probability of getting bruised during sieving. The present study shows that variations in the extraction protocol can alter the total density and community composition of extracted nematodes and provides recommendations for an efficient and standardized approach in future studies. Having a simple, cheap, and standardized extraction protocol can facilitate the assessment of soil biodiversity in global contexts.

## Introduction

Nematodes appear in nearly any kind of soil from dry desert sand to the tundra [1]. Their ubiquitousness, high species richness, and characteristic responses to environmental stressors make them unique biological indicators. Next to information about soil conditions, soil health, and soil processes [2], nematodes and indices based on their community composition can be used to describe soil food web complexity, nutrient enrichment, and decomposition channels (Ferris 2001). These variables can provide important information in ecological research, e.g., studying the consequences of environmental change and biodiversity loss [3–5]. To study general patterns and ecological principles, global networks of ecological experiments have been set up, such as Nutrient Network (https://www.nutnet.umn.edu/), investigating the consequences of multiple nutrient additions in grasslands [6], or TreeDivNet (http://www.treedivnet.ugent.be/) comprising different tree diversity experiments across the globe [7]. Thus far, soil nematodes have mostly been studied in single and local experiments [e.g., 5,8–10], but global assessments of the responses of soil nematodes as bioindicators to environmental changes are scarce. Generally, the assessment of soil biodiversity is largely neglected, leading to a strong under-representation of soil biodiversity in databases, especially at the global scale [11]. Reasons may be a bias towards charismatic species (e.g. vertebrates) and difficulties in sampling procedures.

The high trophic and functional diversity of nematodes comes along with a large number of extraction methods available (reviewed in [12]), highlighting that there may not be one ideal technique for all taxa, and different research questions can ask for different approaches [13]. In addition, the diversity of extraction methods is usually accompanied with complex equipment like the Oostenbrink elutriator [14]. However, labs agreeing to extract nematodes in the frame of global ecological networks need one extraction technique, which can be easily implemented and is low cost.

In this study, we focus on the very common Baermann funnel method [15] as a simple, fast, and cheap approach for nematode extraction in global assessments (Fig 1). By the use of the Baermann funnel method, many samples (hundreds) can be extracted in parallel, and it can be rebuild easily by laboratories without having experience in nematode extraction. The amount of soil needed is relatively small and only a low amount of water is required. Another advantage to other methods is the cleanliness of the final solution (less soil particles) making microscope work easier and faster. However, this method only selects active nematodes, thereby excluding cysts and inactive forms.

**Fig 1.**
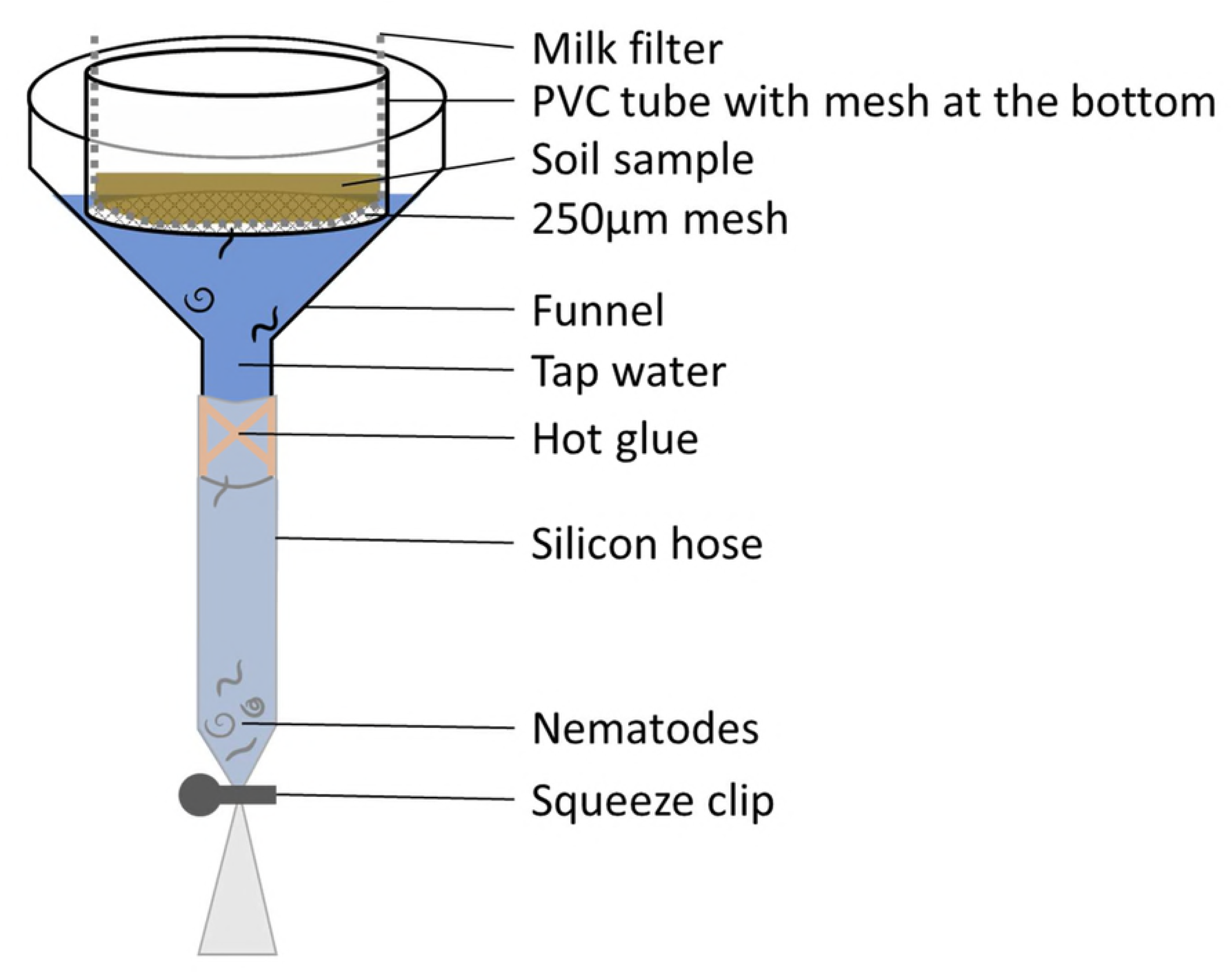
Schematic Baermann funnel apparatus for nematode extraction from soil. A silicone hose is fixed with hot glue to a funnel with an inner diameter of 11 cm. The end of the funnel is closed with a clip to prevent leaking of the nematode solution. A PVC tube of an inner diameter of 7 cm with a 250 μm mesh at the bottom is covered with a milk filter to prevent soil particles to enter the soil solution, which is than filled with soil. The apparatus is filled with tap water until it touches the soil (no submerging) to moisten the soil and increase nematode movement. Nematodes will accumulate at the bottom of the closed silicone hose.

Building on many previous studies testing nematode extraction efficiencies [13,16,17], we provide a detailed protocol to extract soil nematodes, e.g., for global assessments, focusing on a consistent methodology and high extraction efficiency, which can be used by many laboratories worldwide. By doing so, we highlight how common soil pre-treatments and equipment (sieving, amount of soil, filter type) can affect extraction efficiency and community composition and should therefore be considered in future studies.

Generally, soil samples are homogenized before extraction by sieving [18]. Mesh sizes from 1 mm up to 5 mm are commonly used, whereby small mesh sizes require gentle sieving to not bruise nematodes [12]. Mortality can occur when samples are handled roughly, and loss of species after sieving was shown to be species-specific [13,19]. By contrast, homogenization breaks soil aggregates and is assumed to facilitate the movement of nematodes through the soil, likely resulting in higher nematode extraction efficiency. However, despite the inconsistency of mesh sizes used in different studies, the consequences for extraction efficiency and comparability of results have not been tested before.

In ecological long-term experiments or in pristine habitats, destructive samplings like taking soil cores often are strongly limited to prevent destruction of the plots, and only small amounts of soil may be available for nematode extraction. Using large amounts of soil, on the other hand, may also reduce extraction efficiency as less mobile nematodes are discriminated [12,17]. Thus, exploring the role of the amount of extracted soil for nematode extraction efficiency is required to provide general recommendations.

Different permeable filters are used to separate nematodes from soil. Most often cotton-wool milk filters are used, but also cheesecloth, filter paper, or paper tissue are suggested. However, knowledge of the influence of different filters on extraction efficiency is missing. Regarding the availability of materials and costs, we test milk filters and common paper towels in this study.

A well-chosen combination of the settings described above may help to increase nematode extraction efficiency and to avoid potential biases of different extraction protocols. In this study, we evaluated different settings of the Baermann-funnel method by varying 1) different sieving mesh sizes, 2) different amounts of soil, and 3) two different filter types to investigate the consequences for the total amount of extracted nematodes and for nematode community composition. In addition, two very different soil types, i.e., loamy and sandy soil, were used to enable us to make general recommendations.

## Methods

We tested the effects of four variables in soil nematode extraction in a full-factorial design: two soil types (loamy and sandy soil), three soil sieving treatments (2 mm mesh size, 5 mm mesh size, no sieving), three amounts of extracted soil (25 g, 50 g, and 100 g fresh weight), and two permeable filters (milk filters and paper towels). All treatments were replicated five times resulting in 180 samples.

The loamy soil was taken from the Jena Experiment, a grassland biodiversity experiment in Jena, Germany [20]. Adjacent to the experimental plots, soil samples were taken from a soil depth of 0 to 20 cm with pH 8.1, carbon concentration 4.6%, nitrogen concentration 0.3%, and C-to-N ratio 15.7. Clay content was 14%, silt content 41%, and sand content 45% [21]. The sandy soil was taken from the Kreinitz Experiment, a tree biodiversity experiment in Zeithain, Saxony, Germany [22]. Soil samples were taken at a distance of around 10 m to the experimental plots from 0 to 20 cm depth. Soil pH was 5.5, carbon concentration 1.1 %, nitrogen concentration 0.1%, and C-to-N ratio was 11.4. Clay content was 2%, silt content 5%, and sand 94%.

Before any treatments were applied, soil was gently mixed. Afterwards, a fraction of the soil was sieved with a mesh of 2 mm or 5 mm. One fraction of the soil was not sieved, but roots and stones were removed by hand to correctly evaluate soil weight [12]. Three different amounts of fresh soil were used for extraction: 25 g, 50 g, and 100 g, representing a thickness of the soil during extraction of about 1, 2, and 4 cm, respectively. Finally, two different filter types were used: commonly used milk filters (Sana, type FT 25) and paper towels (ZVG Zellstoff-Vertriebs-GmbH & Co. KG, EAN: 4026899028532).

### Baermann funnel apparatus

The Baermann funnel apparatus consisted of a funnel with an inner diameter of 11 cm. Using a different diameter is possible but should be constant among and within studies. A 12 cm piece of silicone hose was attached to the funnel ending and fixed with hot glue to prevent water leakage. The tube was closed with a squeezer clip at the end of the silicon hose. The Baermann funnel apparatus was installed in a horizontal position without buckling of the silicon hose. Soil of a certain amount was filled into circular PVC tubes (7 cm in diameter) with a mesh of 250 μm at the bottom, allowing nematodes to traverse the mesh. The mesh was covered with a filter (milk filter or paper towel) to prevent soil particles to enter the nematode solution (Fig 1). To obtain clean samples, we used a large piece of the filter material to prevent soil particles to enter the nematode solution from the side. This, however, increases evaporation and water has to be added if necessary to prevent that the soil falls dry.

### Procedure

The first step was to check if the apparatus is tight by filling fresh tap water (room temperature or below) into the funnel with the closed silicone hose at the bottom until it reached the lower end of the funnel. The silicone hose had to be squeezed several times to remove air from the silicone hose.

The weight of the empty PVC tube including the filter and a label to identify the sample was noted. Fresh soil of 25 g, 50 g, or 100 g was filled into the PVC tubes. The exact weight has to be noted to get soil water content and relate nematodes to g dry soil. The height of the soil volume was 0.9 cm, 1.9 cm, and 3.8 cm for 25 g, 50 g, and 100 g of fresh soil weight, respectively. Afterwards, the PVC tube with soil was inserted in the funnel.

Fresh tap water was added from the side until the bottom of the mesh of the PVC tube touched the water to saturate the sample with water to increase nematode mobility. Samples were not submerged with water to prevent oxygen limitation. After 72 h [12], nematodes accumulating at the bottom of the tube were transferred into vials and fixed in 4% hot formalin. Therefore, the silicone hose was opened and the water containing nematodes rinsed through a sieve with a 15 μm mesh to separate nematodes from water. Nematodes accumulating on the mesh were transferred into a vial by rinsing the mesh with hot formalin (4%). After extraction, the soil with the PVC tube were dried and weighed to obtain nematode densities per g soil dry weight.

### Nematode counting and identification

All nematodes within one sample were counted with a microscope (Leica, DMI 3000 B) using 50× magnification. To detect changes in nematode community composition due to the different treatments, a subset of treatments showing the strongest differences in the total number of extracted nematodes (see below) were identified to family level after Bongers (1994) and Andrássy (2005) using 1000× magnification. We randomly identified 100 individuals per sample. The proportional value of each family was extrapolated to the total number of nematodes in the sample.

### Statistical analysis

The full-factorial experiment was analyzed using four-way Analysis of Variance (ANOVA) with the factors soil sieving (three levels: 5 mm mesh size, 2 mm mesh size, no sieving), amount of extracted soil (three levels: 25 g, 50 g, and 100 g fresh soil), filter type (two levels: milk filters and paper towels), and all possible interactions. Each treatment was replicated five times resulting in 180 samples. As four-way interactions are complex, we simplified the analysis by separating the dataset by soil type as main differences arouse due to strong differences in densities (mean ± sd of nematodes extracted from 1 g of sandy soil was 1.4 ±0.9 compared to samples from the loamy soil with 17.5 ± 12.4, respectively.) Analyses were performed using R (R i386 3.3.1; [25]). Model residuals were checked and fulfilled assumptions of the performed tests.

We used principal component analysis (PCA) to detect if specific nematode families were selected by the different extraction treatments. Based on the strongest treatments effects on the total number of extracted nematodes, three treatments with n = 3 were chosen for more detailed identification and PCA analysis: *i*) 100 g soil fresh weight sieved at 5 mm, *ii*) 25 g soil fresh weight sieved at 5 mm, and *iii*) 25 g soil fresh weight without sieving. All samples used for the multivariate analysis were extracted with milk filter. As we assumed nematode body size to reflect different levels of mobility, three size classes were considered, i.e., small (up to 0.5 mm), intermediate (0.5 to 1.0 mm), and large (>1.0 mm). Therefore, the mean size of all species/genera per family was calculated (Table 1) using values listed in Bongers (1994), and in Andrássy (2005) for the family Microlaimidae. In addition, nematode families were assigned to the five c-p classes according to [26] and [27] reflecting life strategie histories with cp 1 and cp 2 indicate r-strategists and cp 3 to cp 5 indicate K-strategists. Furthermore, nematode families were classified according their occurrence to abundant (up to 5%), medium (5-1%), and rare (below 1%) families using mean relative occurrence of nematode families in loamy and sandy soils. PCA was performed with R i386 3.3.1 [25].

**Table 1.**
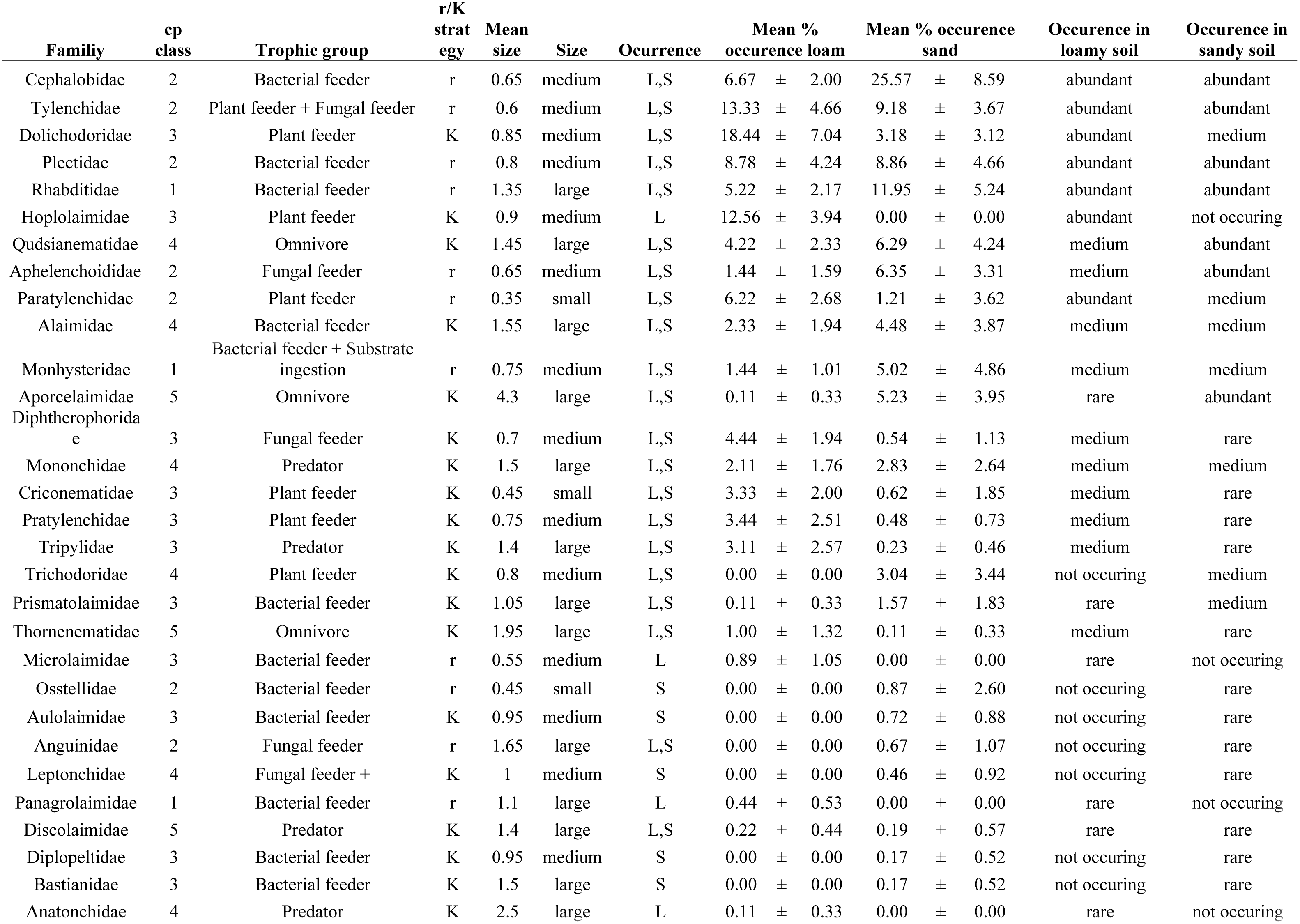
Nematode families extracted from loamy and sandy soil. List of nematode families extracted from loamy (L) and sandy (S) soil using the Baermann funnel method with assigned c-p classes after Bongers (1990) and Bongers and Bongers (1998) and trophic groups after Yeates et al., 1993). In addition, assignment to r- and K strategists. R-strategsits were nematodes from c-p class 1 and 2, whereas c-p classes 3 to 5 are classified as K-strategists [36]. Size classes (small, intermediate, and large) of all nematode families were calculated as the mean size of the minimum (min) and maximum (max) of all genera and species belonging to one family being listed in Bongers (1994), and in Andrássy (2005) for the family Microlaimidae. Occurrence describes in which soil types nematodes occurred. Using mean relative occurrence of nematode families in loamy and sandy soils were used to assign nematodes to abundant (up to 5%), medium (5-1%), and rare (below 1%) families. Means ± SD of nematode families are given for samples displayed in the PCA using different extraction treatments, i.e., using *i*) 25 g of fresh soil sieved at 5 mm and *ii*) no sieving, respectively, as well as using 100 g of fresh soil sieved at 5 mm (n = 3). Taxa were sorted by overall mean of extracted nematodes.

| Family | cp<br>class | Trophic group | r/K<br>strat<br>egy | Mean<br>size | Size | Ocurrence | Mean %<br>occurrence loam |  | Mean % occurrence<br>sand |  | Occurrence in<br>loamy soil | Occurrence in<br>sandy soil |
| --- | --- | --- | --- | --- | --- | --- | --- | --- | --- | --- | --- | --- |
| Cephalobidae | 2 | Bacterial feeder | r | 0.65 | medium | L,S | 6.67 | ± 2.00 | 25.57 | ± 8.59 | abundant | abundant |
| Tylenchidae | 2 | Plant feeder + Fungal feeder | r | 0.6 | medium | L,S | 13.33 | ± 4.66 | 9.18 | ± 3.67 | abundant | abundant |
| Dolichodoridae | 3 | Plant feeder | K | 0.85 | medium | L,S | 18.44 | ± 7.04 | 3.18 | ± 3.12 | abundant | medium |
| Plectidae | 2 | Bacterial feeder | r | 0.8 | medium | L,S | 8.78 | ± 4.24 | 8.86 | ± 4.66 | abundant | abundant |
| Rhabditidae | 1 | Bacterial feeder | r | 1.35 | large | L,S | 5.22 | ± 2.17 | 11.95 | ± 5.24 | abundant | abundant |
| Hoplolaimidae | 3 | Plant feeder | K | 0.9 | medium | L | 12.56 | ± 3.94 | 0.00 | ± 0.00 | abundant | not occurring |
| Qudsianematidae | 4 | Omnivore | K | 1.45 | large | L,S | 4.22 | ± 2.33 | 6.29 | ± 4.24 | medium | abundant |
| Aphelenchoididae | 2 | Fungal feeder | r | 0.65 | medium | L,S | 1.44 | ± 1.59 | 6.35 | ± 3.31 | medium | abundant |
| Paratylenchidae | 2 | Plant feeder | r | 0.35 | small | L,S | 6.22 | ± 2.68 | 1.21 | ± 3.62 | abundant | medium |
| Alaimidae | 4 | Bacterial feeder | K | 1.55 | large | L,S | 2.33 | ± 1.94 | 4.48 | ± 3.87 | medium | medium |

|  |  |  |  |  |  |  |  |  |  |  |  |  |  |  |
| --- | --- | --- | --- | --- | --- | --- | --- | --- | --- | --- | --- | --- | --- | --- |
| Monhysteridae | 1 | Bacterial feeder + Substrate ingestion | r | 0.75 | medium | L,S | 1.44 | ± | 1.01 | 5.02 | ± | 4.86 | medium | medium |
| Aporcelaimidae | 5 | Omnivore | K | 4.3 | large | L,S | 0.11 | ± | 0.33 | 5.23 | ± | 3.95 | rare | abundant |
| Diphtherophoridae | 3 | Fungal feeder | K | 0.7 | medium | L,S | 4.44 | ± | 1.94 | 0.54 | ± | 1.13 | medium | rare |
| Mononchidae | 4 | Predator | K | 1.5 | large | L,S | 2.11 | ± | 1.76 | 2.83 | ± | 2.64 | medium | medium |
| Criconematidae | 3 | Plant feeder | K | 0.45 | small | L,S | 3.33 | ± | 2.00 | 0.62 | ± | 1.85 | medium | rare |
| Pratylenchidae | 3 | Plant feeder | K | 0.75 | medium | L,S | 3.44 | ± | 2.51 | 0.48 | ± | 0.73 | medium | rare |
| Tripylidae | 3 | Predator | K | 1.4 | large | L,S | 3.11 | ± | 2.57 | 0.23 | ± | 0.46 | medium | rare |
| Trichodoridae | 4 | Plant feeder | K | 0.8 | medium | L,S | 0.00 | ± | 0.00 | 3.04 | ± | 3.44 | not occurring | medium |
| Prismatolaimidae | 3 | Bacterial feeder | K | 1.05 | large | L,S | 0.11 | ± | 0.33 | 1.57 | ± | 1.83 | rare | medium |
| Thornenematidae | 5 | Omnivore | K | 1.95 | large | L,S | 1.00 | ± | 1.32 | 0.11 | ± | 0.33 | medium | rare |
| Microlaimidae | 3 | Bacterial feeder | r | 0.55 | medium | L | 0.89 | ± | 1.05 | 0.00 | ± | 0.00 | rare | not occurring |
| Ostellidae | 2 | Bacterial feeder | r | 0.45 | small | S | 0.00 | ± | 0.00 | 0.87 | ± | 2.60 | not occurring | rare |
| Aulolaimidae | 3 | Bacterial feeder | K | 0.95 | medium | S | 0.00 | ± | 0.00 | 0.72 | ± | 0.88 | not occurring | rare |
| Anguinidae | 2 | Fungal feeder | r | 1.65 | large | L,S | 0.00 | ± | 0.00 | 0.67 | ± | 1.07 | not occurring | rare |
| Leptonchidae | 4 | Fungal feeder + | K | 1 | medium | S | 0.00 | ± | 0.00 | 0.46 | ± | 0.92 | not occurring | rare |
| Panagrolaimidae | 1 | Bacterial feeder | r | 1.1 | large | L | 0.44 | ± | 0.53 | 0.00 | ± | 0.00 | rare | not occurring |
| Discolaimidae | 5 | Predator | K | 1.4 | large | L,S | 0.22 | ± | 0.44 | 0.19 | ± | 0.57 | rare | rare |
| Diplopeltidae | 3 | Bacterial feeder | K | 0.95 | medium | S | 0.00 | ± | 0.00 | 0.17 | ± | 0.52 | not occurring | rare |
| Bastianidae | 3 | Bacterial feeder | K | 1.5 | large | S | 0.00 | ± | 0.00 | 0.17 | ± | 0.52 | not occurring | rare |
| Anatonchidae | 4 | Predator | K | 2.5 | large | L | 0.11 | ± | 0.33 | 0.00 | ± | 0.00 | rare | not occurring |

## Results

Nematode extraction efficiency was affected by a significant four-way interaction of soil type, sieving, amount of soil, and filter (Table S1). To better identify specific treatment effects, datasets were divided by soil type as nematode densities were mainly affected by soil type. Mean ± sd nematode density in loamy soil was 17.5 ± 12.4 nematodes g^−1^ dry soil compared to 1.4 ± 0.9 nematodes g^−1^ dry soil in sandy soil. The combination of treatments was of importance for nematode extraction efficiency in the loamy soil as indicated by the three-way interaction, but this was not the case in the sandy soil (Table 2).

**Table 2.**
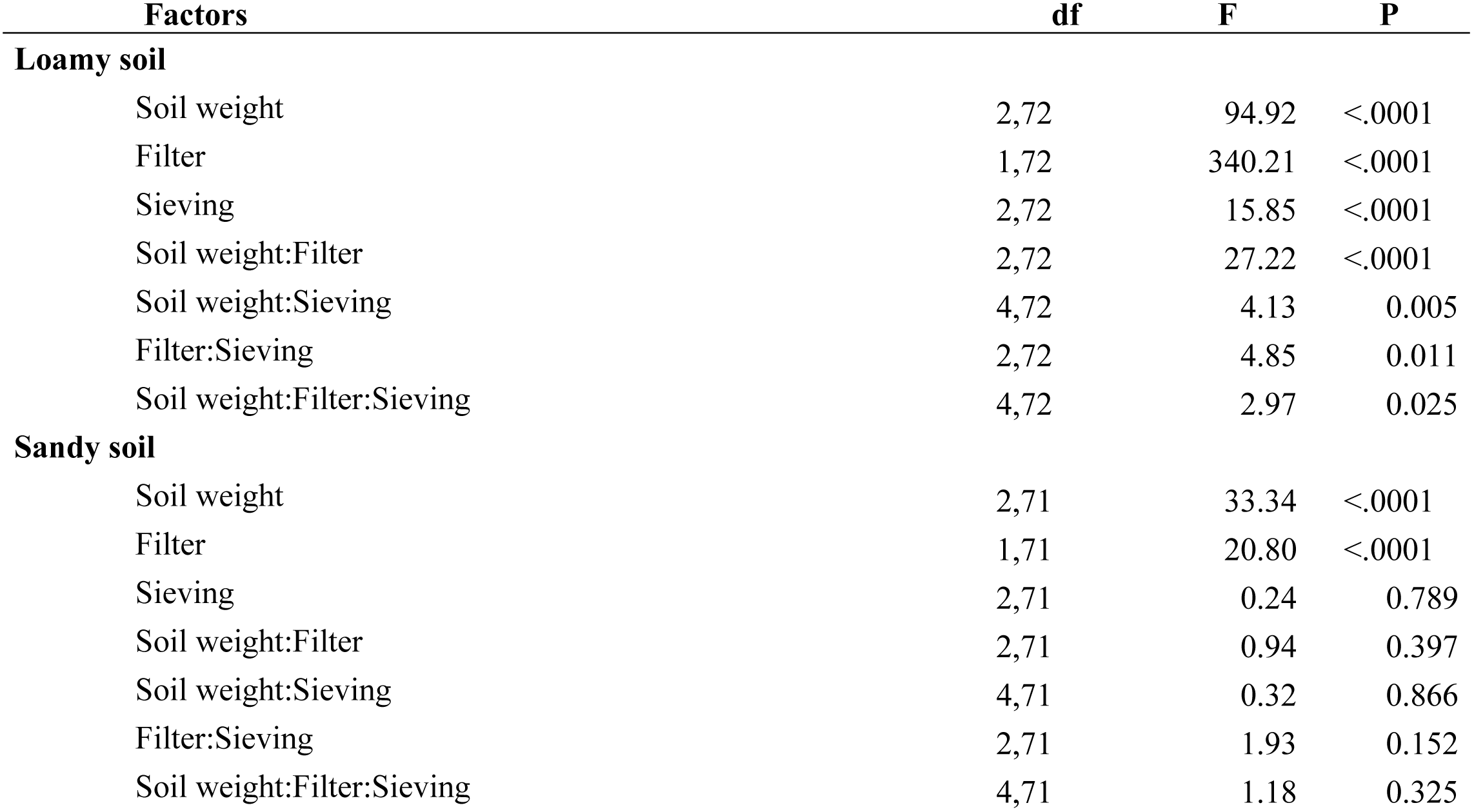
Treatment effects on nematode extraction. ANOVA table of F and P values of the effect of soil weight (25 g, 50 g, and 100 g fresh soil), type of filter (milk filter and paper towel), and sieving (2 mm, 5mm, no sieving), and all possible interactions on nematode extraction efficiency (total nematode densities expressed as individuals g^−1^ dry soil) in two different soil types (loamy and sandy soil) using the Baermann funnel technique. df: degrees of freedom. Significant results are marked in bold.

In loamy soil, highest nematode extraction efficiency was achieved using 25 g of fresh soil sieved with a mesh size of 5 mm and using milk filters (42.2 ± 7.7 nematodes g^−1^ dry soil weight), with milk filters generally increasing nematode extraction efficiency (Fig 2a). Sieving 25 g of soil with a smaller mesh resulted in only slightly fewer nematodes (40.4 ± 5.6 nematodes g^−1^ dry soil weight; -4% in comparison to sieving at 5 mm) and did not differ significantly from sieving with 5 mm. In contrast, nematode extraction efficiency was significantly lower when the soil was not sieved. No sieving of 25 g fresh soil resulted in 31.8 nematodes g dry soil^−1^, that is, 25% fewer nematodes compared to highest number of extracted nematodes. In sandy soil, sieving was not of significant importance (Table 2, Fig 2b). Similar as for the loamy soil though, increasing the amount of soil decreased nematode extraction efficiency significantly (Table 2; Fig 2c).

**Fig 2.**
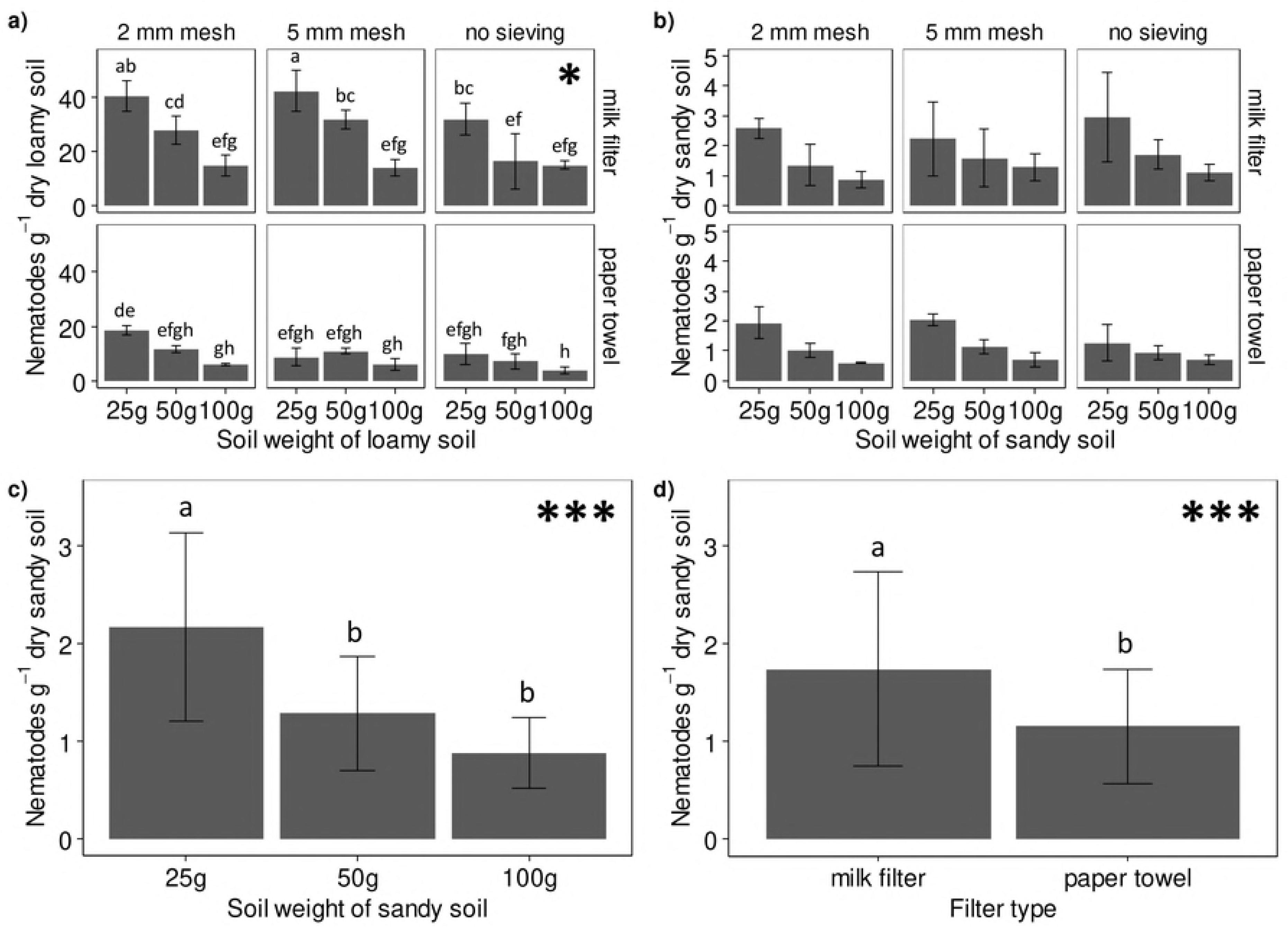
Treatment effects on nematode extraction efficiency. Nematode extraction efficiency of the Baermann funnel method in a) loamy and b-d) sandy soil as affected by sieving soil with different mesh sizes (2 mm, 5mm, no sieving), using different filters (milk filter and paper towels) to obtain clean samples, and by using different amounts of fresh soil (25 g, 50 g, and 100 g of fresh soil weight). Significant single factor effects of the sandy soil are given in c) for soil weight and in d) for filter type as indicated by asterisks. Asterisk in panel a) shows the significant three way interaction. Different letters indicate significant difference (Tukey’s HSD; α=0.05). ^*\**^P *< 0.05, ^***^*P *< 0.001.*

Generally, paper towels significantly reduced overall nematode efficiency by 65% in loamy soil and by 34% in sandy soil. Increasing the amount of soil decreased nematode extraction efficiency from 25 g to 50 g by 30% and from 25 g to 100 by 61% in loamy soil, whereas in sandy soil the reduction was 42% and 60%, respectively (Table 3).

**Table 3.**
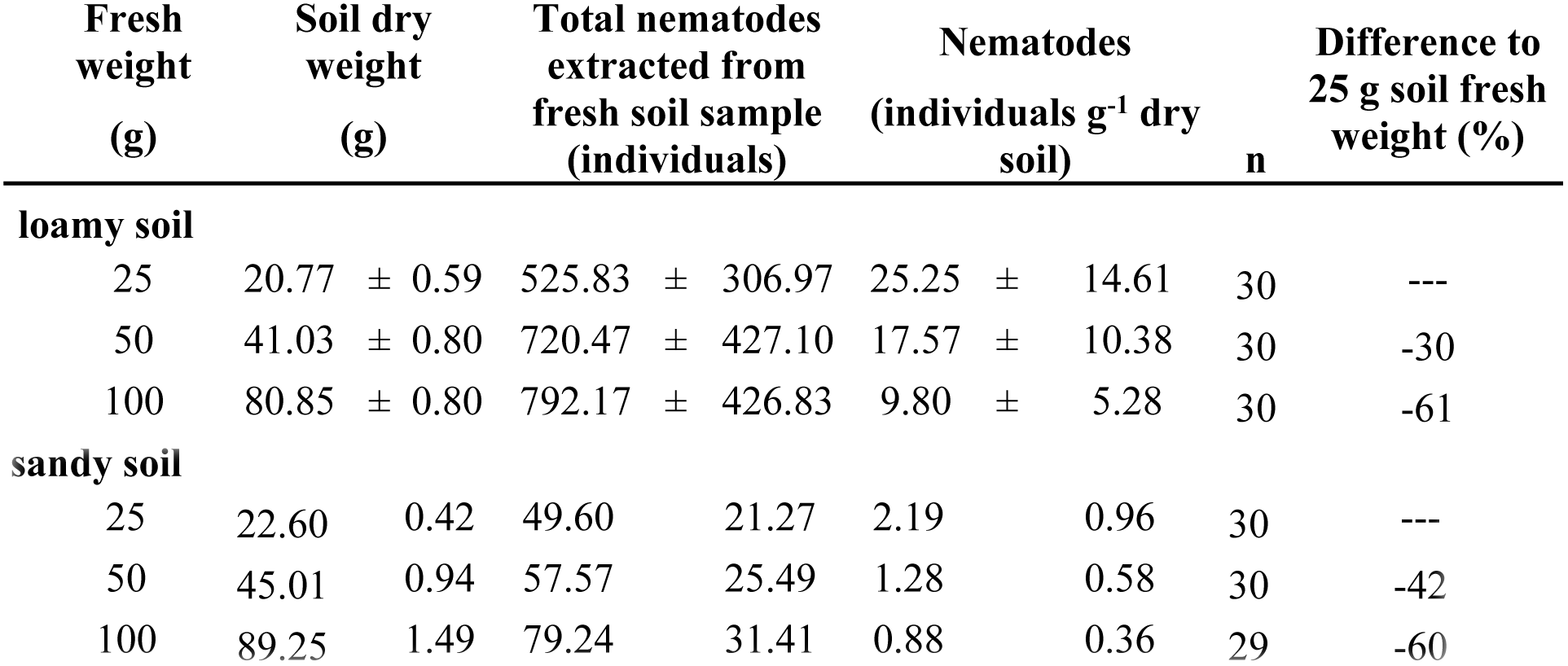
Comparison of nematode extraction efficiency and total amount of extracted nematodes. Extracted nematodes from three different amounts of fresh soil (25 g, 50 g, and 100 g) related to fresh and dry soil weight and the proportional difference between soil amounts.

| Fresh weight<br>(g) | Soil dry weight<br>(g) | Total nematodes extracted from<br>fresh soil sample<br>(individuals) | Nematodes<br>(individuals g<sup>-1</sup> dry<br>soil) | n | Difference to<br>25 g soil fresh<br>weight (%) |
| --- | --- | --- | --- | --- | --- |
| **loamy soil** |  |  |  |  |  |
| 25 | 20.77 ± 0.59 | 525.83 ± 306.97 | 25.25 ± 14.61 | 30 | --- |
| 50 | 41.03 ± 0.80 | 720.47 ± 427.10 | 17.57 ± 10.38 | 30 | -30 |
| 100 | 80.85 ± 0.80 | 792.17 ± 426.83 | 9.80 ± 5.28 | 30 | -61 |

**sandy soil**
|  |  |  |  |  |  |  |  |  |
| --- | --- | --- | --- | --- | --- | --- | --- | --- |
| 25 | 22.60 | 0.42 | 49.60 | 21.27 | 2.19 | 0.96 | 30 | --- |
| 50 | 45.01 | 0.94 | 57.57 | 25.49 | 1.28 | 0.58 | 30 | -42 |
| 100 | 89.25 | 1.49 | 79.24 | 31.41 | 0.88 | 0.36 | 29 | -60 |

To assess the consequences of sieving and soil weight on nematode community composition, we used a selection of three specific treatment combinations (all using milk filters as densities were highest here), i.e., *i*) sieving with 5 mm and using 100 g fresh soil (lowest yield in loamy soil), *ii*) sieving with 5 mm and using 25 g of fresh soil (highest yield in loamy soil), and *iii*) no sieving and using 25 g of fresh soil (medium yield in loamy soil). Generally, nematode family composition differed strongly between soil types/sites (explained 58.6% of the variation, first axis; Fig 3). In the loamy soil, extraction treatments had a stronger effect of the family composition, whereas in sandy soil family composition was more homogenous and sieving and soil weight were of lower importance. In loamy soil, family composition in samples extracted from 100 g fresh soil was more different from samples using 25 g of fresh soil, whereas the family composition of samples without sieved soil and using 25 g fresh soil was intermediate. In loamy soil, more K-strategists were extracted from 100 g fresh soil compared to 25 g fresh soil (Fig 4a). No sieving did not increase the amount of K-strategists, i.e., larger organisms that are supposed to be more likely to be damaged by sieving (Fig S1). However, in samples using 100 g loamy soil, more large (Fig S1) and rare (Fig S2) nematodes were extracted. Generally, nematode families from different c-p classes distributed more homogenously among the treatments in sandy soils, indicating a less strong effect of sieving and soil weight in sandy soils.

**Fig 3.**
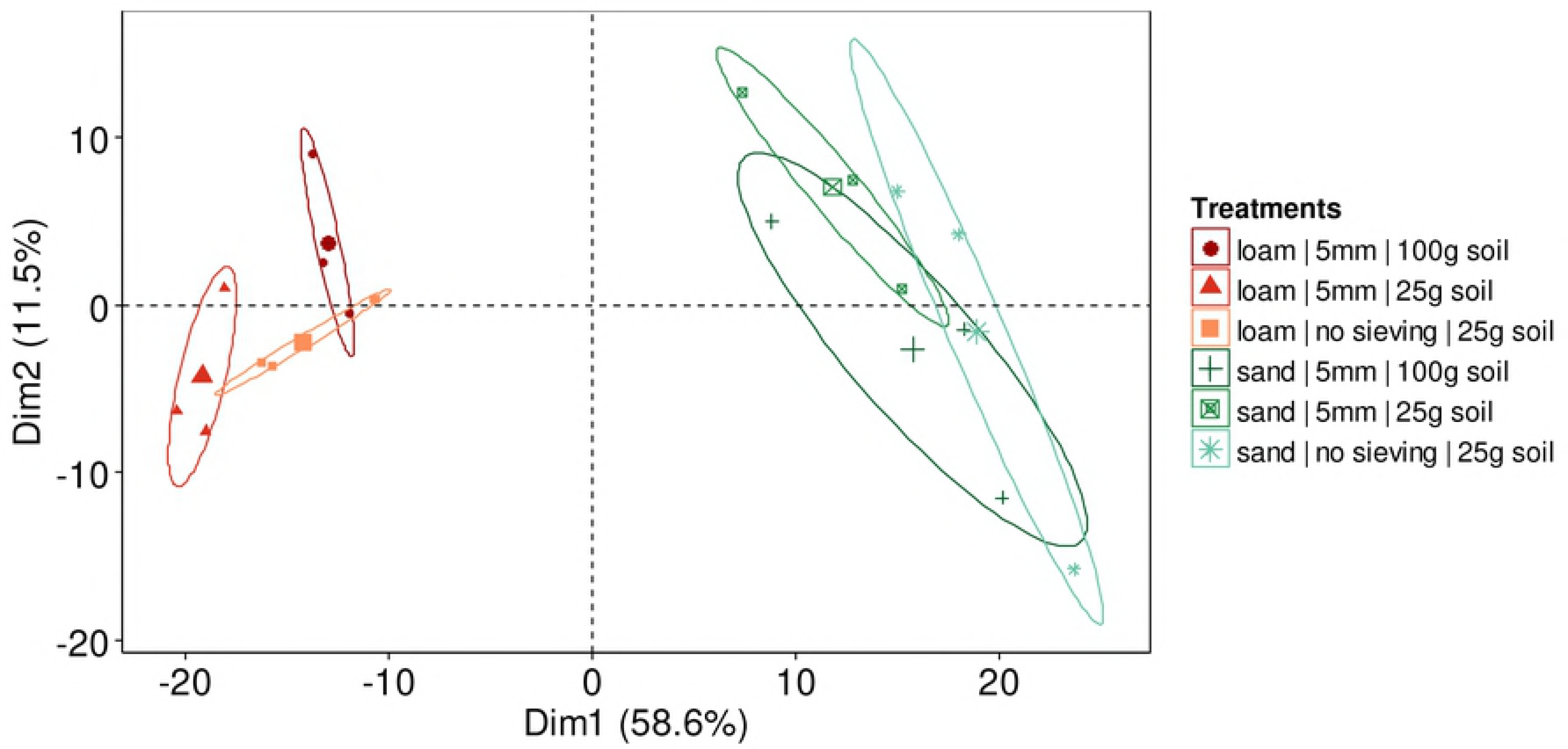
Effect of soil type and soil pre-treatments on nematode community composition. Principal component analysis (PCA) of the nematode community (family level) as affected by soil type and different treatments (sieving with 5 mm mesh size and no sieving, and different amounts of soil [25 g and 100g fresh soil]) prior to extraction reflecting treatment combinations with highest (sieving with 5 mm and 25 g soil), medium (no sieving and 25 g soil), and lowest (5 mm sieving and 100 g soil) nematode extraction efficiency in loamy soil. Symbols represent the specific treatment combinations with larger symbols display centroids. Numbers in brackets are variation explained by the first (Dim1) and second (Dim2) PCA axis, respectively.

**Fig 4.**
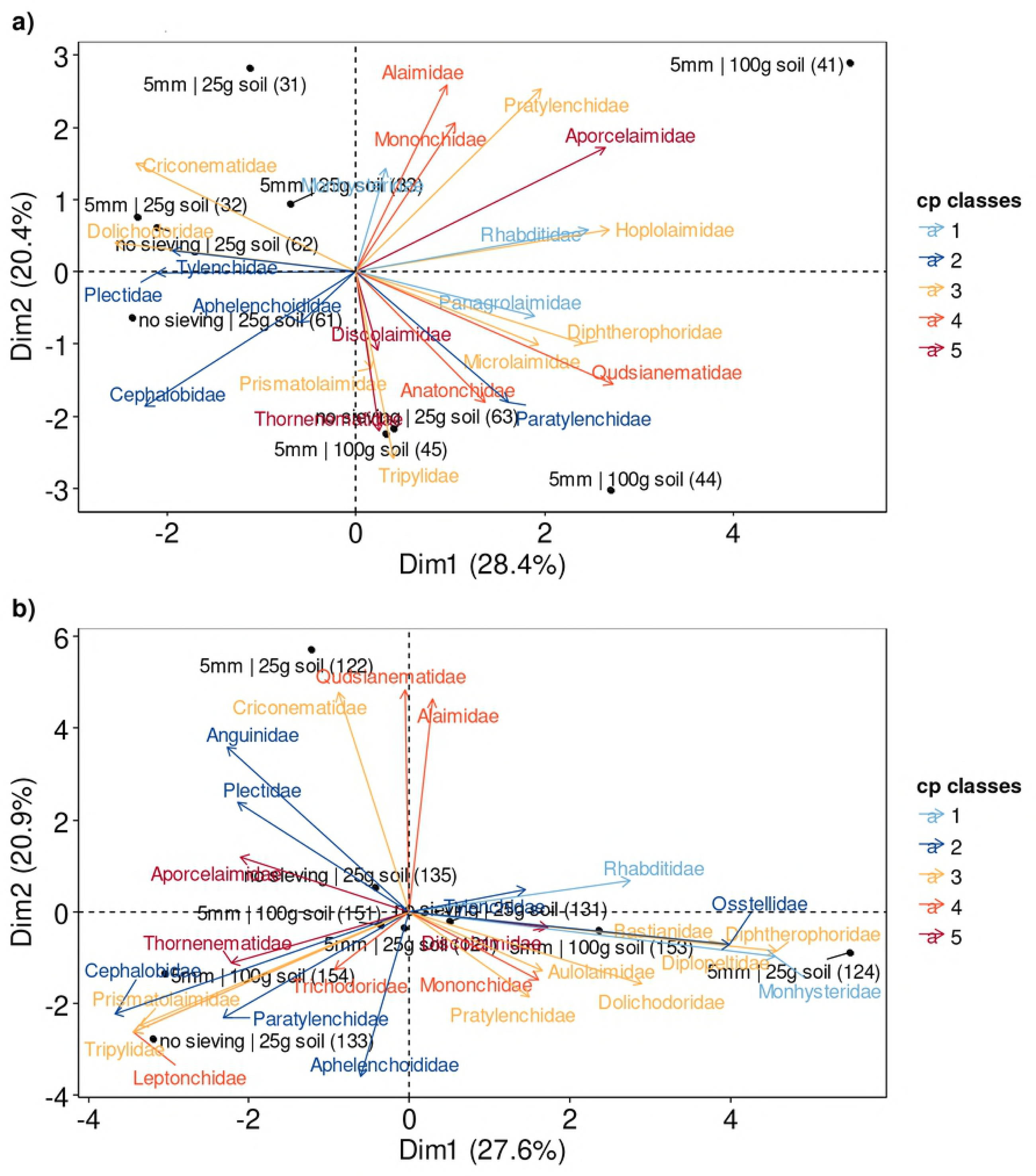
Distribution of nematode c-p classes after extracting nematode with different pre-treatments. Principal component analysis (PCA) of the nematode community (family level) as affected by different treatments (sieving with 5 mm mesh size and no sieving, and different amounts of soil [25 g and 100g fresh soil]) prior to extraction reflecting treatment combinations with highest (sieving with 5 mm and 25 g soil), medium (no sieving and 25 g soil), and lowest (5 mm sieving and 100 g soil) nematode extraction efficiency in a a) loamy and b) sandy soil. Nematode families were assigned to the five c-p classes according to Bongers (1990) and Bongers & Bongers (1998) with blueish colors indicate r-strategists and reddish colors K-strategists. Numbers in brackets are variation explained by the first (Dim1) and second (Dim2) PCA axis, respectively.

## Discussion

In the present study, we found that the combination of different extraction treatments significantly affected nematode extraction efficiency. Although treatment combinations were of different importance in loamy and sandy soil, overall highest numbers of extracted nematodes were observed when using milk filters and the lowest amount of soil, i.e., 25 g fresh soil. Sieving was important in the loamy soil, while it did not matter if soil was sieved with 2 or 5 mm mesh size. Although not sieving soil yielded highest nematode numbers in the sandy soil, no significant difference was found between sieving and not sieving. Therefore, to achieve high nematode extraction efficiency in different types of soil, it is recommended to use small amounts of soil in combination with sieving and using milk filters.

**Sieving** was of higher importance in the loamy soil than in the sandy soil. Loamy soil has more stable soil aggregates than sandy soil, which is why we suggest that breaking up soil aggregates by sieving increases nematode mobility in loamy soil as nematodes are no longer limited by soil structure and pore space [28]. Using 5 mm compared to 2 mm mesh size resulted in slightly higher (4%) extraction efficiency and may reflect losing some rare families when using a smaller mesh size. These rare families were larger in body size in the present study and may have had a higher probability to be injured by sieving with 2 mm. As we did not analyze the community composition of samples sieved with 2 mm, we cannot provide specific information about the consequences for nematode communities. However, as the observed effect was small (4%) and non-significant, we believe that both mesh sizes can be recommended.

Extraction efficiency decreased with an increased **amount of soil**. The higher proportion of large nematodes in 100 g loamy soil suggests that small nematodes may not have been able to pass and or exit thick soil volumes during the common extraction time of 72 h. [29] observed an entomopathogenic nematode to overcome maximally 80 mm in 14 days, indicating that nematodes may be rather slow, which is why the standard [12] even suggests to use a soil volume of only a few millimeters in height. In addition, a thick soil layer can reduce oxygen supply [30], which may decrease nematode survival in the soil sample. PCA revealed that using 100 g of soil increased the number of extracted nematodes of rare families. This may be the result of a higher probability that 100 g soil contains more rare species. Moreover, rare species often are large in body size, which is why they may have a higher chance to be extracted from larger amounts of soil, as mentioned above. However, the soil volume of 100 g samples also selected against the majority of other nematode families. In summary, using the combination of treatments that resulted in highest nematode extraction efficiency may select against some rare species but may better reflect total densities. To overcome this tradeoff, reducing the thickness of the soil layer by increasing the diameter of the funnel/PVC tube-system may help to increase the amount of soil used and the surface area, allowing nematodes to exit the soil for improved qualitative and quantitative nematode community assessments.

Using **milk filters** resulted in a significantly higher number of extracted nematodes than using paper towels. Paper towels are supposed to adsorb water, whereas milk filters are supposed to filter a solution. The fabric of paper towels is probably chosen such that the fibers will take up water, and this paper structure may hamper nematodes to pass the paper towel. Instead of using paper towels as an alternative for milk filters for biodiversity assessments, they may be used to artificially reduce nematode densities according to morphological traits and alter community composition for targeted experiments.

Although we analyzed only a small fraction of possible treatment combinations on the family level, we were able to show that pretreating the soil can change the community composition of extracted nematodes. These results highlight the need to standardize nematode extraction protocols and to account for potential differences when comparing data from multiple sites and studies in syntheses and meta-analyses. The present study may guide the implementation of common nematode extraction protocols for future research.

Nematodes are a powerful indicator taxon, and global assessments of soil nematode communities could increase our understanding of global distribution patterns. Generally, only few datasets of global belowground biodiversity exist [31–34], but these still have insufficient data coverage. In this study, we present a rather simple method, i.e. the Baermann funnel method, to extract nematodes from different soils. The simplicity of the method and the suggested standardized approach allows also non-experts to extract nematodes to participate in global soil biodiversity assessments.

## Acknowledgements

We thank UFZ to access the Kreinitz tree diversity platform and two anonymous reviewers for previous comments, which strongly improved the manuscript. In addition, we thank Anja Zeuner for extracting nematodes from the sandy soil.

## Supporting information

**S1 Table. Treatment effects on nematode extraction.** ANOVA table of F and P values of the effect of soil type (loamy and sand soil), sieving (2 mm mesh size, 5mm mesh size, no sieving), type of filter (milk filter and paper towel), soil weight (25 g, 50 g, and 100 g fresh soil) and all possible interactions on nematode extraction efficiency (total nematode densities expressed as individuals g^−1^ dry soil) using the Baermann funnel technique. df: degrees of freedom.

**S1 Fig. Distribution of nematode size classes after extracting nematode with different pre-treatments.** Principal component analysis (PCA) of the nematode community (family level) as affected by different treatments (sieving with 5 mm mesh size and no sieving, and different amounts of soil [25 g and 100g fresh soil]) prior to extraction reflecting treatment combinations with highest (sieving with 5 mm and 25 g soil), medium (no sieving and 25 g soil), and lowest (5 mm sieving and 100 g soil) nematode extraction efficiency in a a) loamy and b) sandy soil. Nematode families were assigned to different size classes according to overall mean values of individuals calculated from sizes given in Bongers (1994) and Andrássy (2005). Numbers in brackets are variation explained by the first (Dim1) and second (Dim2) PCA axis, respectively.

**S2 Fig. Distribution of nematode occurrence after extracting nematode with different pre-treatments.** Principal component analysis (PCA) of the nematode community (family level) as affected by different treatments (sieving with 5 mm mesh size and no sieving, and different amounts of soil [25 g and 100g fresh soil]) prior to extraction reflecting treatment combinations with highest (sieving with 5 mm and 25 g soil), medium (no sieving and 25 g soil), and lowest (5 mm sieving and 100 g soil) nematode extraction efficiency in a a) loamy and b) sandy soil. Nematode families were assigned according their mean relative occurrence to abundant (up to 5%), medium (5-1%), and rare (below 1%). Numbers in brackets are variation explained by the first (Dim1) and second (Dim2) PCA axis, respectively.

**S1 File. Raw data of total numbers of nematodes extracted.** 180 samples were taken in a full four-way factorial design with the factor soil type (loam and sand), sieving (2 and 5 mm mesh size, no sieving), filter (milk filter and paper towels), and soil weight (25 g, 50 g and 100 g fresh soil weight). Total numbers of extracted nematodes are provided as well as nematodes related to g dry soil.

**S2 File. Raw data of nematode family composition.** Identified nematode families from a subset (n=15) of samples for estimating nematode extraction efficiency. Selected treatments had highest, lowest and medium nematode extraction efficiency in loamy soil.

